# Dual-site tACS over the primary motor cortices increases interhemispheric inhibition and improves bimanual dexterity: A triple-blind, randomised, sham-controlled study

**DOI:** 10.1101/2024.10.27.620547

**Authors:** Brooke Lebihan, Lauren Mobers, Shannae Daley, Ruth Battle, Natasia Leclercq, Katherine Misic, Kym Wansbrough, Ann-Maree Vallence, Alexander D. Tang, Michael A. Nitsche, Hakuei Fujiyama

## Abstract

Concurrent application of transcranial alternating current stimulation (tACS) over distant cortical regions has been shown to modulate functional connectivity between stimulated regions; however, the precise mechanisms remain unclear. Here, we investigated how dual-site tACS (ds-tACS) applied over the bilateral primary motor cortices (M1s) modulates connectivity between M1s. Using a cross-over sham-controlled triple-blind within-subject design, 37 (27 female, age 18-37yrs) healthy participants received tACS (1.0mA, 20Hz) over the bilateral M1s for 20 min. Before and after tACS, functional connectivity between M1s was assessed using imaginary coherence (ImCoh) measured via resting-state electroencephalography (EEG) and interhemispheric inhibition (IHI) via dual-site transcranial magnetic stimulation (TMS) protocol. Additionally, manual dexterity was assessed using the Purdue pegboard task. While ImCoh remained unchanged after simulation, spectral power analysis showed a significant decrease in beta (20 Hz) power during the tACS session. ds-tACS but not sham strengthened IHI between the M1s and improved bimanual assembly performance. These results suggest that improvement in bimanual performance may be explained by modulation in M1-M1 IHI, rather than by coupling in the oscillatory activity. As functional connectivity underlies many clinical symptoms in neurological and psychiatric disorders, these findings are invaluable in developing non-invasive therapeutic interventions that target neural networks to alleviate symptoms.

---

Transcranial alternating current stimulation (tACS) employs weak, alternating currents and is thought to have neuromodulatory effects by inducing phase synchronisation between endogenous brain oscillations and the applied tACS frequency (Thut et al. 2012; Herrmann et al. 2013). The simultaneous application of tACS to distant cortical sites has been shown to alter communication between the stimulated regions, indicating a modulatory effect on functional connectivity (Polanía et al. 2012). Furthermore, it is believed that tACS promotes long-term potentiation-like plasticity even after the stimulation ends, suggesting its potential to induce enduring changes in brain connectivity and function (Vossen et al. 2015; Wischnewski et al. 2019). Impaired functional connectivity underlies various psychiatric, neurological and age-related conditions (O’Reilly et al. 2017; Marzola et al. 2023; Zhang et al. 2023; Castellanos et al.). Therefore, enhancing functional connectivity between cortical sites positions tACS as a powerful tool for research with potential therapeutic applications (Tavakoli & Yun 2017).

Modulating functional connectivity by simultaneously stimulating distant cortical regions by tACS is based on established principles of brain network dynamics. The ‘communication-by-coherence’ hypothesis posits that neuronal communication is optimal when the neural oscillatory phases of the communicating cortices are synchronised (Fries 2005; Fries 2015). Given that motor and cognitive tasks are not executed by a single brain region in isolation but involve complex interactions across multiple brain regions (e.g., Battleday et al. 2014), it is reasonable to assume that simultaneous stimulation of functionally relevant cortical regions via tACS may result in improved functional communication between the stimulated regions (Violante et al. 2017; Reinhart & Nguyen 2019; Grover et al. 2022; Meng et al. 2023).

Polanía and colleagues (2012) empirically demonstrated that dual-site tACS (ds-tACS) modulated functional connectivity between cortical regions during a working memory task. They applied 0° relative phase (in-phase) and 180° relative phase (anti-phase) stimulation to the left prefrontal and parietal cortices, finding that in-phase stimulation enhanced working memory performance while anti-phase stimulation led to a performance decline. These results suggest that tACS likely modified the phase alignment of endogenous oscillatory activity within fronto-parietal networks that are instrumental in working memory performance. Further research by Fujiyama and colleagues (2023) showed that in-phase beta frequency (20 Hz) ds-tACS applied over the right inferior frontal gyrus and pre-supplementary motor area improved response inhibition and increased task-related functional connectivity observed with electroencephalography (EEG). Similarly, other studies have reported beneficial effects of ds-tACS on various cognitive and motor functions, particularly in the theta and beta frequency ranges (Violante et al. 2017; Reinhart & Nguyen 2019; e.g., Grover et al. 2022; Meng et al. 2023). While these findings demonstrate that ds-tACS can enhance functional connectivity and related behaviours, the exact mechanisms underlying these effects remain elusive. To further elucidate these mechanisms, a multi-modal approach combining different neurophysiological techniques could be beneficial.

EEG studies suggest that tACS induces oscillation entrainment, synchronising brain activities with the phase of the applied tACS frequency (Thut et al. 2012; Herrmann et al. 2013). In-vivo animal studies indicate that applied sinusoidal currents entrain endogenous oscillations (Reato et al. 2013); however, evidence remains suggestive but unclear in humans. Evidence of oscillatory entrainment primarily focuses on how tACS induces effects during stimulation. However, behavioural changes that persist beyond the stimulation period are thought to result from plasticity mechanisms (Polanía et al. 2012). Specifically, the ongoing synchronisation of oscillatory activity by tACS appears to regulate the precise timing of neural spikes, leading to long-term potentiation demonstrably distinct from entrainment-related aftereffects (Vossen et al. 2015). Alternate theories suggest that plasticity mechanisms may better explain how tACS induces oscillatory changes and subsequent behavioural changes (Vossen et al. 2015). While EEG has a high temporal resolution, which is beneficial for characterising oscillatory coupling induced by ds-tACS, it provides relatively limited spatial resolution, offering a global measure of neural activity with less spatial specificity in characterising the dual-site tACS-induced changes in cortical connectivity. In contrast, transcranial magnetic stimulation (TMS) offers greater spatial precision, particularly when investigating connectivity between primary motor cortices (M1s). Dual-site TMS (ds-TMS) with paired electromyography (EMG) protocols can be used to investigate M1-M1 interhemispheric inhibition (IHI) (e.g., Bäumer et al. 2006; Koch et al. 2006; Mochizuki et al. 2004; Ni et al. 2009; O’Shea et al. 2007), facilitating the precise assessment of intracortical M1 circuits and the connected contralateral corticospinal neurons, enabling a more precise understanding of the interactions between cortical regions (Ferbert et al. 1992).

This study aimed to investigate the neurophysiological mechanisms underlying tACS-induced changes in functional connectivity between the bilateral M1s. A multimodal approach was utilised employing EEG, ds-TMS, and a behavioural measure using the Purdue pegboard to examine the effect of ds-tACS and sham on neuronal phase synchronisation, IHI, and manual dexterity, respectively. We specifically focus on interhemispheric interactions between the M1s to answer respective research questions since the assessment protocols for M1-M1 functional and structural connectivity and its involvement in motor functions are well-established (e.g., Bäumer et al. 2006; Fujiyama et al. 2016; Ni et al. 2009). We predict that the simultaneous application of beta-frequency in-phase tACS would enhance functional connectivity between the M1 regions, observed across neurophysiological measures, and would consequently result in improved manual dexterity when compared to sham. Identifying the underlying mechanisms by which tACS improves functional connectivity is crucial, as enhancing functional connectivity holds significant clinical potential for developing future interventions for both health and disease.

## Materials and methods

### Participants

A power analysis was performed for sample size estimation using the “simr” package (Lenth 2020) in R statistical package, version 4.4.1 (R Core Team 2024), drawing on data from our previous studies investigating the effect of non-invasive brain stimulation on neurophysiological measures (Fujiyama et al. 2016; 2022). With an alpha level set at 0.05 and a target power of 0.85, consistent with recommendations for minimising the risk of Type II errors in neurophysiological studies (Lakens 2013), we estimated that a sample size of 34 participants would be sufficient to observe a large pre-post change (Cohen’s *d* = 0.8), as supported by previous studies with similar designs (e.g., Fujiyama 2016; 2022) employing neurophysiological measures. To ensure a conservative estimate and enhance statistical power, we recruited 37 (27 female) participants, which provides a power of approximately 0.9.

Participants were aged 18-37 years (*M* = 24.70 years, *SD* = 6.18 years) and were recruited through a university research participation portal. Student participants earned testing time-equivalent course credits for both the real-tACS and sham sessions or received an AUD20 gift voucher following completion of the final session. All participants were screened for non-invasive brain stimulation contraindications, and those with psychiatric or neurological conditions or prescription medications that conflicted with stimulation were excluded (Rossi et al. 2021). The Edinburgh Handedness Inventory (Oldfield 1971) was used to assess participant handedness. Only those with a score above 40, indicating right-handedness, were eligible to participate, as left-handedness has been associated with differing motor cortical representations and non-invasive simulation aftereffects (Nicolini et al. 2019; Fitzgerald et al. 2021). Written informed consent was obtained before participation in the study. This study was approved by the Murdoch University Ethics Committee (2021/240).

## Materials

### Transcranial alternating current stimulation (tACS)

ds-tACS was delivered with a neuroConn DC-STIMULATOR MC (NeuroConn, Ilmenau, Germany) using round rubber electrodes and conductive ten20 paste (Weaver and Company, Aurora, CO, USA) in a 4×1 montage (Figure 1A). A central electrode (2 cm diameter) was positioned over the scalp site targeting the hand representation of M1 in each hemisphere, identified by TMS (see *3.2.2. Transcranial magnetic stimulation (TMS) and electromyography (EMG) recording* section for more detail), with four surrounding return electrodes (2 cm diameter) placed in a radius of 3 cm, measured from the centre of the central electrode to the centre of each return electrode(Villamar et al. 2013), which is the optimal configuration for M1 focal stimulation (Edwards et al. 2013). Alternating currents were applied at 20Hz at 1.0mA peak-to-peak amplitude and with zero DC offset to both stimulation sites at 0° phase lag.

**Fig. 1.**
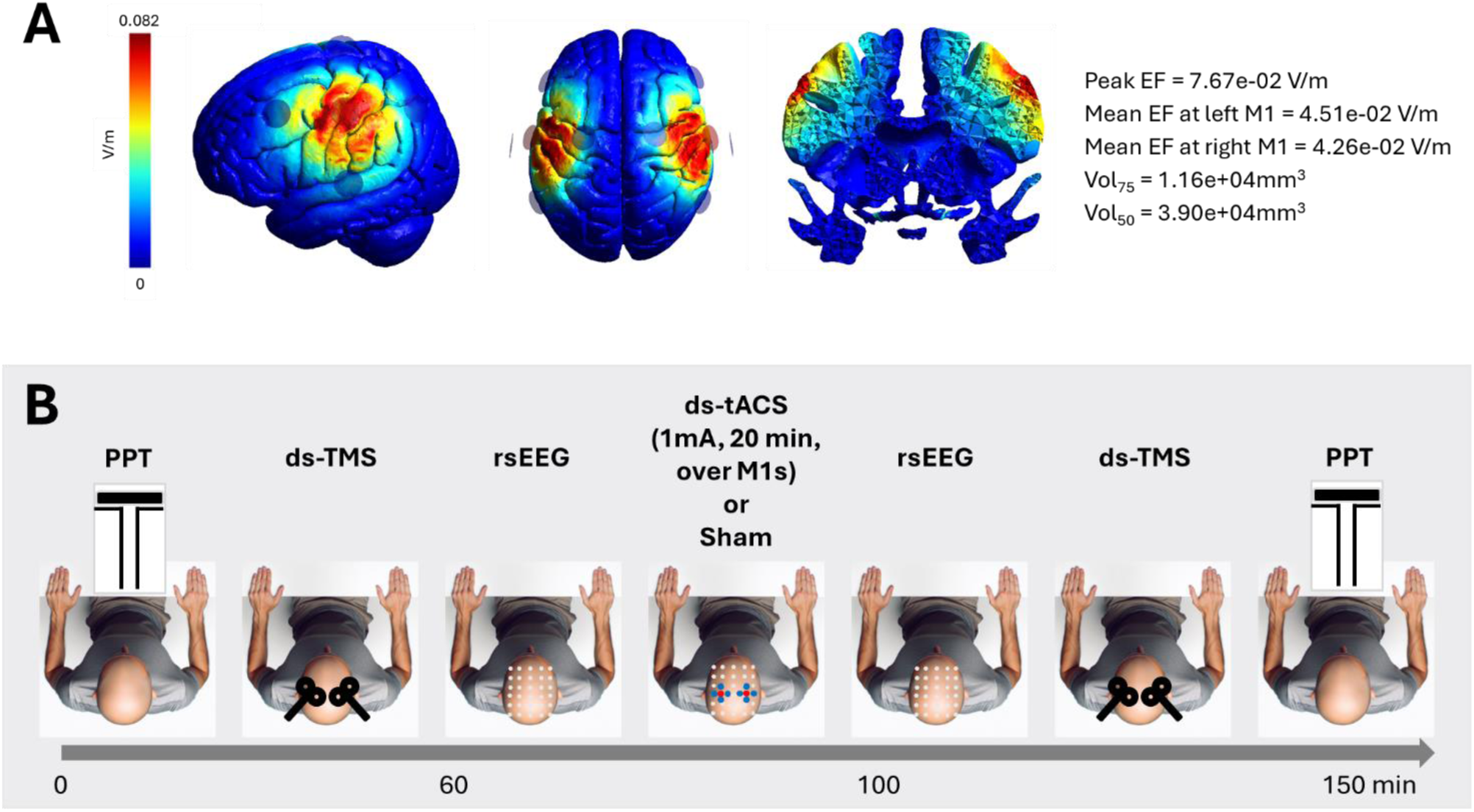
**A.** Stimulated putative norm electric field modelling for tACS electrode montage over the M1s (left and middle panels). A 4 x 1 electrode montage was used for each M1. Currents with peak-to-peak amplitudes of 1mA were delivered to the stimulation sites with a 0° phase differences. The current flow simulation was conducted via SimNiBS 4.0 (Santurino et al., 2019) on the MNI152 head model. Peak electric field strength was defined as the 99.9th percentile of the field. MNI coordinates of the human motor area template (Mayka et al., 2007) were used to estimate the mean electric field strength at left M1 (−37, −21, 58; radius 10 mm) and right M1 (37, −21, 58; radius 10 mm). Vol50 and Vol75: mesh volume with field strength ≥ 50% of the 99.9th percentile and ≥ 75% of the 99.9th percentile, respectively. **B.** Schematic representation of the procedural order of tasks and timeline during tACS and sham and sessions. Each participant attended one tACS and one sham session. PPT = Purdue Pegboard Task, ds-TMS = dual-site transcranial magnetic stimulation, rsEEG=resting state electroencephalography, ds-tACS = dual-site transcranial alternating current stimulation. *Note*: A human figure was generated using Microsoft Copilot.

Two conditions were employed for 21 minutes: tACS at 1.0mA and sham stimulation. During tACS, the intensity ramped up to the target current intensity for 30 seconds and maintained intensity for 20 min before ramping back down at the last 30 seconds. In the sham condition, the tACS stimulation ramped up to 1.0mA and immediately ramped down at the beginning and end of the 20 minutes for 30 seconds to simulate stimulation (Woods et al. 2016). The tACS machine was pre-programmed by a research associate with codes representing the two conditions to ensure both participant and researcher blinding.

### Transcranial magnetic stimulation (TMS) and electromyography (EMG) recording

TMS assessments were performed using two Magstim 200^2^ units (Magstim Company, Dyfed, UK) with two 50mm (D50a coils, Magstim Company, Dyfed, UK), figure-of-eight coils. Surface EMG electrodes recorded TMS-induced motor-evoked potentials from participants’ first dorsal interosseous (FDI) muscles. EMG surface electrodes (Ag/AgCl) were positioned in a belly-tendon montage over the FDI. Signals were amplified with a gain of 1000, band pass filtered (10 - 500 Hz) and sampled at 2000 Hz using a 16-bit AD system (CED1902, Cambridge, UK) for analysis. TMS coils were positioned tangentially, 45° from the sagittal midline, to induce a posterior-anterior current flow, optimal for eliciting MEPs in the FDI, i.e., motor hotspot (Jensen et al. 2005). The motor hotspot was identified by systematically repositioning the coil above the estimated target area until the location that provoked the strongest MEP was found. At the beginning of each session, each individual’s resting motor threshold (rMT) was determined as the lowest intensity that evoked MEPs in the FDI of greater than 50μV in at least three out of five consecutive trials (Carroll et al. 2008) at the marked hotspot.

IHI was assessed using a ds-TMS paradigm following the methodology outlined by Ferbert and colleagues (1992). We investigated M1-M1 projections in both directions (left to right hemisphere and vice versa). The first condition of our protocol involved delivering a single pulse testing stimulus (TS) set at an intensity of 130% of rMT. The following condition employed ds-tMS delivering a conditioning stimulation (CS), intensity set at 110% of rMT, to one M1 followed by the TS to the contralateral M1 at an interstimulus interval (ISI) of 10ms, and then the third condition repeated this double pulse stimulation protocol with an ISI of 40ms (Ni et al. 2009; Kroeger et al. 2010; Hinder et al. 2012). Previous research suggested that short ISIs (≤10 ms) are associated with IHI mediated by postsynaptic GABA_A_ mechanisms, while longer ISIs (≥40 ms) involve GABA_B_-ergic circuits (Sanger et al. 2001; Irlbacher et al. 2007).

At each TMS assessment time point, there were two blocks of IHI, testing left M1- right M1 and right M1- left M1 IHI were conducted in counterbalanced order across sessions and participants. Each block lasted 3 minutes and consisted of 60 trials, alternating between 3 conditions: 20 single-pulse stimuli (TS), 20 CS-TS IHI10 (CS followed 10ms later by TS), and 20 CS-TS IHI40 (CS followed 40ms later by TS) stimuli. The TMS pulses administered throughout hotspot identification, rMT determination, and IHI were applied every 4-6 seconds to avoid inducing neuroplastic changes in the brain (Rossi et al. 2021).

### Electroencephalography

EEG was recorded using a 128-electrode EEG HydroCel Geodesic Sensor Net (Magstim EGI, Eugene, OR). Net Station (5.4.2) software recorded sensor-level EEG signals from Ag-AgCl scalp electrodes. EEG was amplified using a Net Amps 300 amplifier, low and high pass filtered (0.1-500 Hz) with a 1000 Hz sample rate. Electrode impedance was kept below 50 kΩ as recommended by the manufacturer (Magstim EGI, Eugene, OR). During resting-state EEG recording, participants viewed a fixation cross presented on a PC screen for 3 minutes.

### Behavioural Assessment

The Purdue Pegboard Test assessed gross motor ability and fine motor dexterity (Tiffin & Asher 1948). We employed three conditions: unimanual (left and right) and a bimanual assembly task. Participants performed all conditions seated at a table and completed practice trials before the assessment. In the unimanual task, participants were instructed to place as many pegs as possible with one hand into a pegboard within 30 seconds. The number of pegs successfully placed was recorded for each hand. For the assembly task, participants were instructed to use both hands to assemble a peg, washer, collar, and washer down the right column of the pegboard. The number of individual pieces assembled on the board within 60 seconds was recorded.

### Control Measures

#### Sleep, Caffeine and Alcohol Questionnaire

The Sleep, Caffeine, and Alcohol (SCA) questionnaire consists of four questions designed to record various factors that could influence tACS effects and EEG results. It measures sleep quality on a scale from 1 (poor) to 10 (excellent), the number of hours slept, and the amount of caffeine (in milligrams) and alcohol (in units) consumed in the last 12 hours. The SCA was administered during both tACS sessions to account for individual variations that might affect outcomes (Valenzuela 1997; Zulkifly et al. 2021).

#### Transcranial Electrical Stimulation Sensation Questionnaire

After each session, participants completed the Transcranial Electrical Stimulation (tES) Sensation Questionnaire. This questionnaire first assessed 13 common sensations, such as burning, tingling, itching, and headaches, experienced during tACS using a Likert scale ranging from 1 (nothing) to 5 (very strong) (Woods et al. 2016). It also included items to determine when these sensations started (beginning, middle, or end of stimulation) and stopped (soon, in the middle, or at the end of stimulation). Additionally, participants rated how these sensations affected their PPT performance using a 5-point scale from “absolutely not” to “very much.” This questionnaire also evaluated the effectiveness of blinding regarding the condition participants received.

#### Study Design

This study employed a cross-over sham-controlled triple-blind within-subject design to investigate the effect of ds-tACS on M1-M1 functional connectivity and motor function using EEG and TMS. All participants underwent two identical experimental sessions, with assessment measures recorded before and after either real or sham tACS conditions. The order of sessions was counterbalanced across participants, and both sessions occurred at a similar time of day. After the initial session, participants returned for testing a minimum of 7 days later, as the effects of a single session of non-invasive brain stimulation (NiBS) are likely to be diminished by 7 days (e.g., Fujiyama et al. 2017). Following this minimum of 7 days, testing took place at the same time as the primary session to control for the possible effects of cortisol level fluctuation throughout the day that may affect tES responses (Sale et al. 2008).

#### Procedure

Figure 1B illustrates a procedural schematic of the testing sessions. For each session, assessments before the stimulation were carried out in the following order: PPT, TMS, and EEG, while after the stimulation, the same assessments were conducted in reverse order, i.e., EEG, TMS and PPT. The order of the assessments was kept constant across participants for a practical reason. Although TMS and EEG are compatible methods, the presence of an EEG net on the participant’s head requires an increase in TMS intensity. Therefore, the EEG net was applied after the TMS pre-assessment and removed following the post-EEG assessment prior to the post-TMS assessment. The EEG nets remained on the participants’ heads during tACS application, as the tACS electrodes could be applied without the need to remove the nets.

### Data Processing and Statistical Analysis

#### TMS data pre-processing

Corticospinal excitability was determined as an average peak-to-peak MEP in the TS trials of the FDI muscles from 20 to 100 ms post-TMS. The IHI ratio was determined as the mean peak-to-peak MEPs in CS-TS trials relative to the mean amplitude of MEP response to the single TS, i.e., IHI=MEP_CS-TS_/MEP_TS_. The IHI ratio is inversely related to interhemispheric inhibition: a decrease in the IHI ratio indicates increased inhibition, whereas an increase in the IHI ratio indicates reduced inhibition. As such, the ‘IHI ratio’ refers to specific numerical values, while ‘IHI’ describes interhemispheric inhibition in the conventional definition. The trials exceeding the root mean square EMG of 10uV during the 40 ms immediately preceding the TMS pulse were excluded (Carson et al. 2004).

#### EEG data pre-processing

EEG data were pre-processed using the EEGLAB toolbox RELAX (the reduction of electroencephalographic Artifacts) (Bailey et al. 2023) through the MATLAB environment (MathWorks, R2020a). RELAX is a data-cleaning pipeline that reduces vascular, ocular and myogenic-induced artifacts to preserve neural signal readings. Using RELAX, the data was downsampled at 500 Hz. Then, a 50 Hz notch filter and a bandpass filter (1-80 Hz) were applied. Second, deficient EEG sensors and noisy time periods were removed. Third, eye blink and muscle movement-induced artifacts were removed using the Multiple Wiener Filtering method. The Wavelet Enhance Independent Component Analysis removed artifacts not detected in the previous method. Last, once the removed EEG sensors and artifacts were interpolated, the data was divided into two-second segments for statistical analysis.

For ImCoh calculation, the time series of each electrode were convolved with complex Morlet wavelets for frequencies between 4 and 90 Hz in 1 Hz increments (87 wavelet frequencies in total). The length of the wavelets started at 3 cycles for the lowest frequency and logarithmically increased as the frequencies increased, such that the length was 13 cycles for the highest frequency. This approach balances temporal and frequency precision (Cohen 1988).To minimise the effects of edge artifacts, analytic signals were obtained from time windows of 400 to 1600 ms (at 20 ms intervals) within each 2000 ms epoch. ImCoh values were computed for each electrode pair within the C3 (left M1) and C4 (right M1) using the following formula:

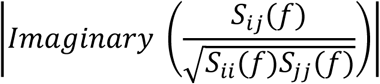

Here, *i* and *j* represented the time series of each electrode. For frequency *f*, the cross-spectral density *S*_*ij*_(*f*) was taken from the complex conjugation of the complex Fourier transforms *x*_*i*_(*f*) and *x*_*j*_(*f*). Coherency was extracted by normalising the cross-spectral density by the square root of the signals’ spectral power, *S*_*ii*_(*f*) and *S*_*jj*_(*f*). Then, the imaginary component of the resultant complex number was obtained. An estimate of M1-M1 ImCoh was obtained by averaging ImCoh values across all electrode pairs at each time point, frequency, and trial.

#### Statistical analysis

For a mixed-effects model, for each dependent variable, within-subject fixed factors of SESSION (tACS/sham) and TIME (pre-stimulation/post-stimulation) were included in the model with by-subject intercept as random effects. For PPT data, an additional fixed factor of CONDITION (left, right, bimanual) was included in the model, while for TMS IHI data, additional factors of ISI (10/40ms) and DIRECTION (LR/RL) were considered in the model. For EEG power and TMS rMT and MEP_TS_ analyses, HEMISPHERE (L/R) was used instead of DIRECTION. A linear mixed effect model (LMM) was used when the data followed a normal distribution curve, while a generalised linear mixed effect model (GLMM) was used when the data deviated from a normal distribution. The assumptions for the G/LMMs, including linearity, homogeneity of variance, and normal distribution of the model’s residuals, were evaluated using the “DHARMa” package (Hartig 2024), which employs a simulation-based method to examine residuals for fitted G/LMMs. Null hypothesis significance testing for main and interaction effects was conducted with Wald Chi-Squared tests for the GLMM analyses and *F-*tests for LMM analyses, and significant main and interaction effects were further investigated with Bonferroni corrected contrasts. Standardised effect sizes were not reported for each fixed term in the G/LMM, following recommendations by Pek & Flora (2018). The partitioning of variance within the G/LMM makes obtaining a standardised effect size for each model term difficult. Instead, to facilitate the interpretation of G/LMM results, Cohen’s *ds* for follow-up contrasts were provided alongside significance levels, the critical *p*-value was set at .05.

For the control measures, the SCA and tES sensation questionnaire, the Shapiro-Wilk test was conducted to assess the normality of the data. For normally distributed variables, paired sample t-tests were conducted. For non-normally distributed variables, Wilcoxon signed-rank tests were conducted.

Statistical analyses and visual illustrations were performed using R statistical package, version 4.4.1 (R Core Team, 2024) with an integrated environment, RStudio version 2024.04.2+764 (RStudio Team, 2020) for Windows using “lmerTest” v3.1-3 (Kuznetsova et al. 2020) to fit LMM and GLMM, “DHARMa” package (Hartig 2024) for LMM assumptions of linearity, homogeneity of variance and normal distribution of residuals “emmeans” v1.10.3 (Lenth 2024) for follow-up contrasts, “ggplot2” 3.5.1 package (Wickham et al. 2016) for graphical plots, “dplyr” v1.1.4 (Wickham et al. 2023) for data manipulation, “janitor” v2.2.0 (Firke 2023) for cleaning data, “here” v1.01 (Muller & Bryan 2020) for declaring paths, “knitr” v1.47 (Xie 2024) for report generation, “reader” v1.0.6 (Cooper 2017) for reading data files, “reshape2” v0.8.1 (Wickham 2007) for reshaping data, “skimr” v2.1.5 (Waring et al. 2022) for data summaries, “stringr” v1.5.1 (Wickham 2023) for string operation wrappers. All data and codes publicly available on the Open Science Framework (OSF): https://osf.io/exf54/.

## Results

During the study, all participants tolerated the interventions without reporting discomfort and no adverse events were experienced during experimental procedures. As shown in Table 1, repeated-measures *t*-tests indicated that participants were likely blinded to the conditions, as there were no significant differences in the transcranial electrical stimulation (tES) sensation questionnaire responses across sessions. Similarly, no significant differences were observed across sessions in the components of the SCA questionnaire.

**Table 1.**
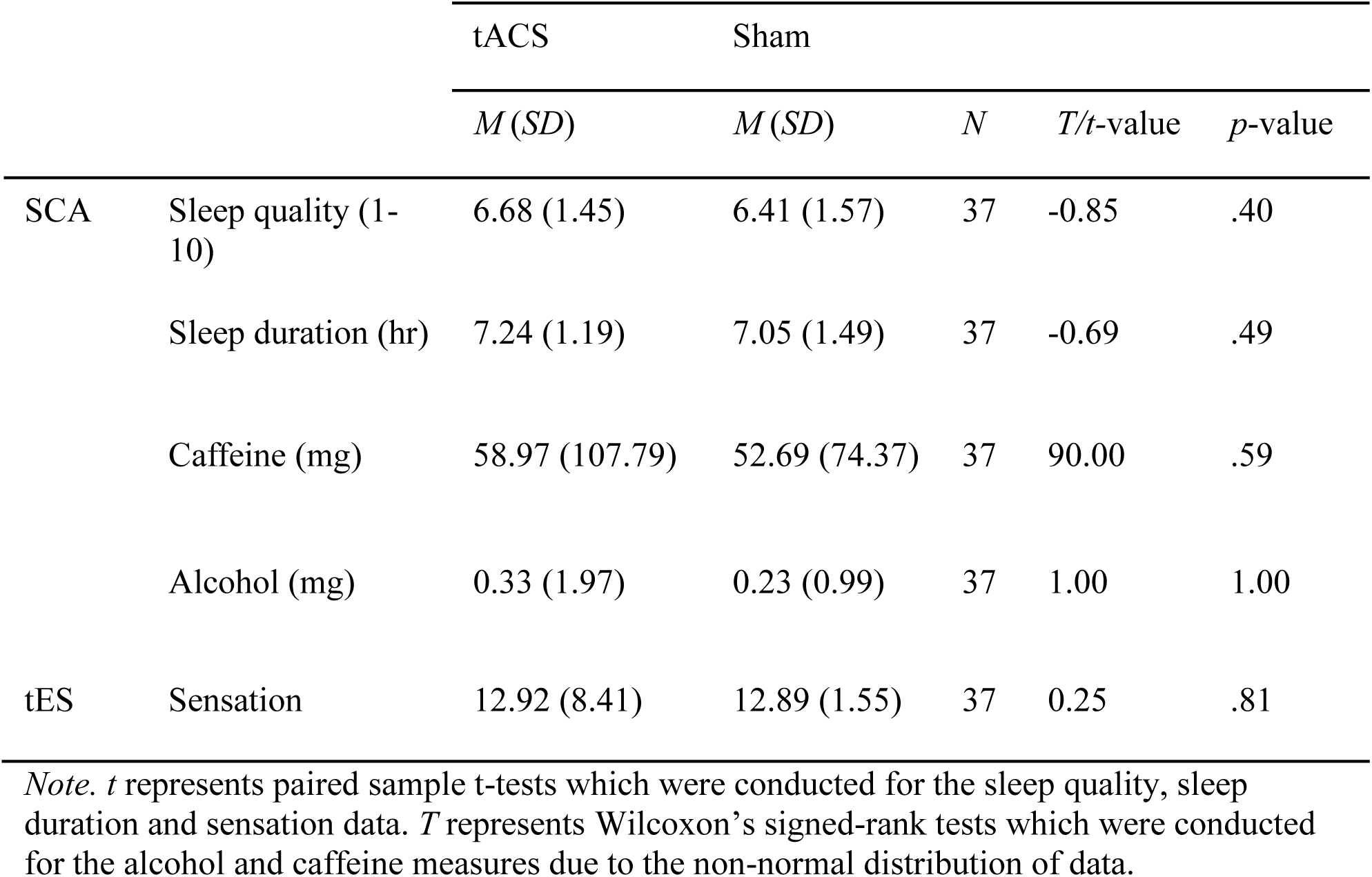
Descriptive statistics and associated t-tests for control measures.

|  |  | tACS | Sham | *N* | *T/t*-value | *p*-value |
| --- | --- | --- | --- | --- | --- | --- |
|  |  | *M* (*SD*) | *M* (*SD*) |  |  |  |
| SCA | Sleep quality (1-10) | 6.68 (1.45) | 6.41 (1.57) | 37 | -0.85 | .40 |
|  | Sleep duration (hr) | 7.24 (1.19) | 7.05 (1.49) | 37 | -0.69 | .49 |
|  | Caffeine (mg) | 58.97 (107.79) | 52.69 (74.37) | 37 | 90.00 | .59 |
|  | Alcohol (mg) | 0.33 (1.97) | 0.23 (0.99) | 37 | 1.00 | 1.00 |
| tES | Sensation | 12.92 (8.41) | 12.89 (1.55) | 37 | 0.25 | .81 |

*Note.* *t* represents paired sample t-tests which were conducted for the sleep quality, sleep duration and sensation data. *T* represents Wilcoxon's signed-rank tests which were conducted for the alcohol and caffeine measures due to the non-normal distribution of data.

### Behavioural Measure

#### Bimanual coordination improves following ds-tACS

Manual dexterity changes following tACS were assessed using three conditions in the PPT: left-hand only, right-hand only, and bimanual assembly. A GLMM revealed a significant main effect of CONDITION, *χ^2^*(2, *N* = 37) = 3939.27, *p* < .001 and a significant interaction of SESSION x TIME x CONDITION *χ^2^*(2, *N* = 37) = 3.34, *p* = .04. Follow up contrasts on the higher order interaction revealed that bimanual dexterity improved significantly by 10.10% (4.32 parts more, i.e., > one assembly) from pre-to-post stimulation in the tACS session, *z* = 3.54, *p* < .01, *d* = 0.90, which was illustrated in Figure 2. This pattern was absent in the sham session, *z* =.05, *p* = .96, *d* = 0.01. Importantly, bimanual assembly performance at post-stimulation was significantly greater in the tACS session than in the sham session, *z* = 2.18, *p* = .03, *d* = 0.56. In contrast, unimanual conditions showed no significant changes across time points for both tACS and sham sessions, |*zs|* = 0.92, *ps* > .36, |*ds|* < 0.23.

**Fig. 2.**
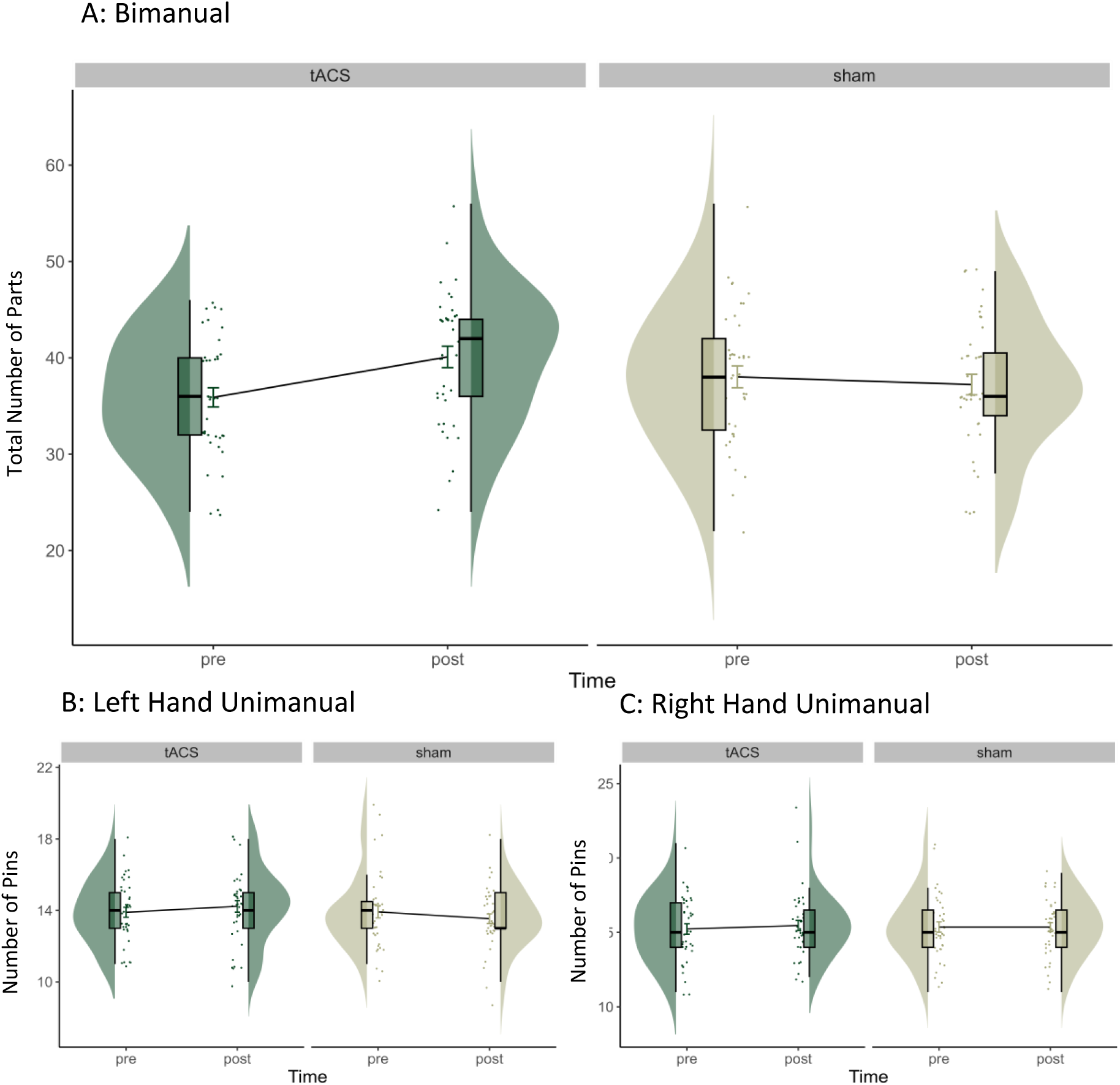
Distribution of pins placed during the Purdue Pegboard Test. This plot compares the effects of tACS and sham stimulation on motor dexterity, measured pre- and post-stimulation, for three task conditions: Bimanual/Both hands assembly (A), Left hand only (B), and Right hand only (C). For the bimanual assembly task, the y-axis represents the total number of parts (pins, collars, washers). The boxplots show the median and interquartile range to visually summarise central tendency and dispersion. The horizontal line within the box plot represents the median. The top and the bottom lines represent the upper and lower quartiles, respectively. The data points outside of the whiskers are >1.5 quartiles. The line connecting the paired violins visualizes the direction of post-stimulation changes.

### Neurophysiological Measures

#### TMS Parameters

Table 2 shows the rMT across sessions for the left and right hemispheres. An LMM revealed that there was a significant main effect of HEMISPHERE, *χ^2^* (1, *N* = 37) = 78.43, *p* < .001. The mean rMT for the right M1 was significantly higher than the left M1 in both sessions. These differences are likely explained by the coils used for each hemisphere. We used the same coil for each M1 across sessions and participants. Due to the stimulators’ position relative to the participant, a longer corded coil was required consistently for the farther side. In addition to the dominant left M1 generally having a lower motor threshold in right-handed participants, the coil used for right M1 stimulation appeared to produce a lower field intensity, leading to a higher rMT than that of the left M1. Since the stimulation intensities for TMS assessments, i.e., single-pulse TMS and IHI, were adjusted based on each individual’s rMT, discrepancies in rMT should not impact the interpretation of TMS results. The main effect of SESSION and the interaction of SESSION x HEMISPHERE, *χ^2^*(1, *N* = 37) = .01, *p* = .91, were not significant, suggesting the differences in rMT between hemispheres were constant across tACS and sham sessions.

**Table 2.** Mean (95 % CI) rMT for the left and right M1 in the tACS and sham sessions.

| rMT |  | tACS | sham |
| --- | --- | --- | --- |
|  | Left M1 | 45.33(8.41) | 45.26(6.54) |
|  | Right M1 | 52.56(8.74) | 52.67(8.02) |

#### Corticospinal excitability decreases following ds-tACS

We investigated how corticospinal excitability changed following ds-tACS to the bilateral M1s. As time-related changes, i.e., pre-tACS vs post-tACS, in corticospinal excitability were of primary interest, only those main effects and interactions involving TIME as a factor are described in detail.

A GLMM revealed significant main effects of TIME, *χ^2^*(1, *N* = 37) = 6.04, *p* = .01 and HEMISPHERE, *χ^2^*(1, *N* = 37) = 25.47, *p* < .01 and significant interactions between TIME x HEMISPHERE *χ^2^*(1, *N* = 37) = 6.67, *p* < .01 and SESSION x HEMISPHERE *χ^2^*(1, *N* = 37) = 6.90, *p* < .01.. The follow up contrasts for the significant interaction of TIME x HEMISPHERE revealed that at baseline, the left M1 (1.79 ± 1.38) showed significantly greater excitability than the right M1 (1.54 ± 1.17), *z* = −5.48, *p* <.001, *d* = −0.246. However, at post, the difference between the left M1 (1.56 ± 1.41) and right M1 (1.44 ± 1.15) became non-significant, *z* = −1.72, *p* =.89, *d* = −0.080. This change was primarily driven by a significant decline in excitability from pre- to post-stimulation in the left M1, *z* = −3.18, *p* = 0.002, *d* = −0.302, while the right M1 showed no significant change, *z* = −1.44, *p* = 0.150, *d* = −.136. These results suggested that, regardless of tACS or sham, the left M1 excitability showed a distinct change over time relative to the right M1 excitability. The follow up contrast for the significant interaction between SESSION x HEMISPHERE revealed that during sham the left M1 exhibited significantly greater corticospinal excitability than the right M1, *z* = −5.52, *p* <.001, *d* = −0.247. During tACS, the left M1 and right M1 corticospinal excitability were not significantly different, *z* = −1.69, *p* =.09, *d* = −0.078. These results indicate that ds-tACS had a more pronounced effect on the left M1, reducing its excitability to a level comparable with the right M1, which remained relatively stable throughout the intervention. The higher-order interactions, including TIME x SESSION, was not significant, *χ^2^s* < 2.92, *ps* > .06. This indicates that the corticospinal excitability change pre-to-post was comparable between tACS and sham sessions, this is illustrated in Figure 3.

**Fig. 3.**
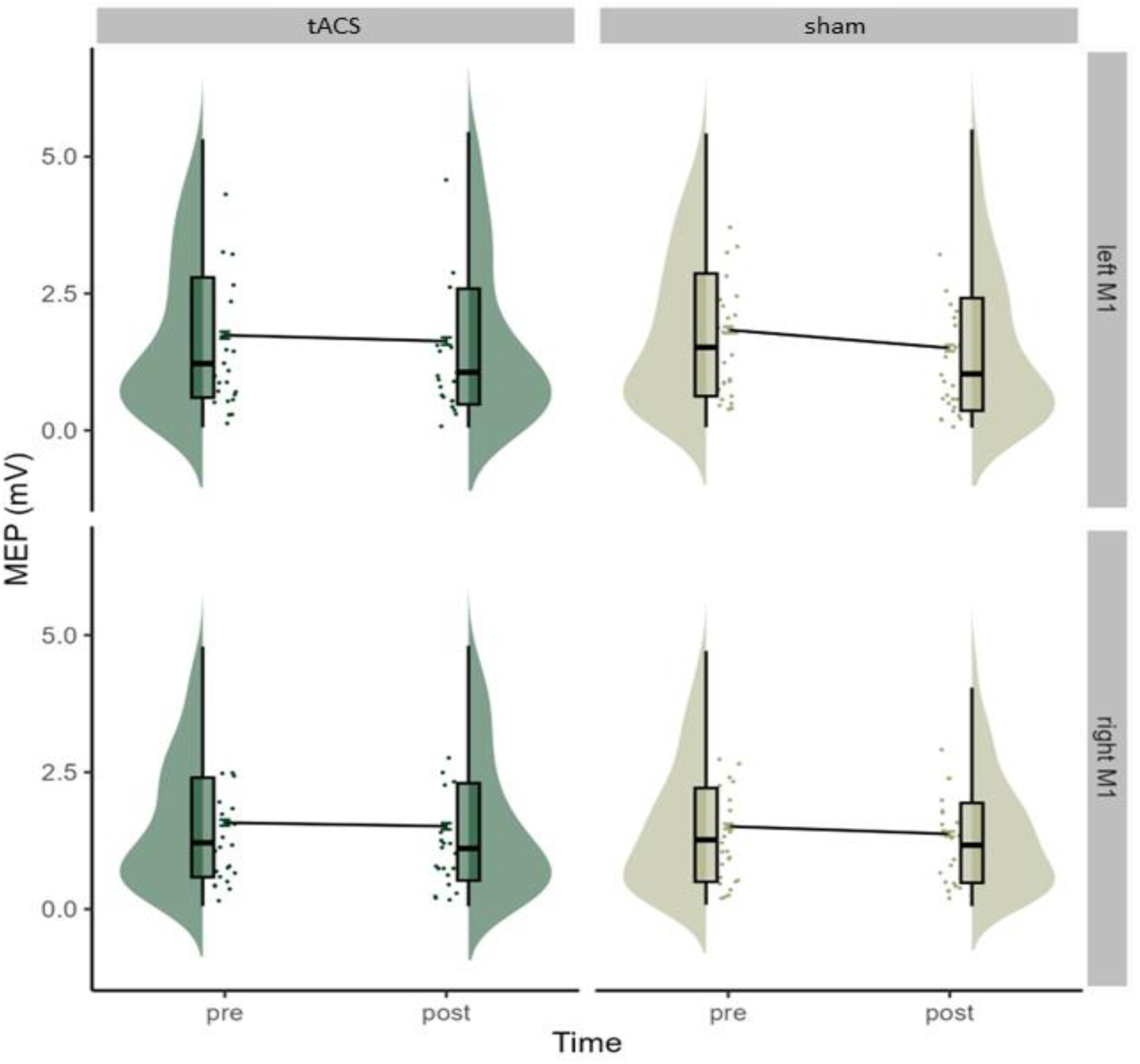
Distributed TS MEP sizes in mV across different conditions: tACS and sham, pre- and post-stimulation. Separate violins represent each condition for both the left and right M1. The width of each violin indicates the density of MEP values. The boxplots show the median and interquartile range to visually summarise central tendency and dispersion. The horizontal line within the box plot represents the median. The top and the bottom lines represent the upper and lower quartile, respectively. The data points outside of the whiskers are >1.5 quartiles. The line connecting the paired violins visualizes the direction of MEP size changes between pre- and post-stimulation.

### IHI is augmented by ds-tACS

A GLMM assessing the IHI ratio revealed a significant main effect of DIRECTION, *χ^2^*(1, *N* = 37) = 11.10, *p* < .01. The IHI ratio was significantly higher for the left to right direction (0.79 ± 0.17) than right to left direction (0.72 ± 0.21), indicating that at rest, the inhibitory projection from the left M1 to the right M1 was less prominent than from right M1 to the left M1. GLMM also revealed a significant interaction of SESSION x TIME, *χ^2^*(1, *N* = 37) = 5.53, *p* = .02. As shown in Figure 4, follow-up contrasts revealed that during the tACS session, the IHI ratio significantly decreased (19.3% reduction) from pre-stimulation to post-stimulation, *z* = −2.40, *p* = .02, *d* = −.92, indicating increased interhemispheric inhibition between M1s, while, in the sham session, IHI ratio did not change significantly, *z* = 0.19, *p*=.85, *d* = .04. In addition, at post-stimulation, the IHI ratio was significantly lower in the tACS session compared to the sham session, *z* = −2.00, *p* = .045, *d* = −.81, while baseline IHI ratios were comparable between the tACS and sham session, *z* = 0.71, *p* = .48, *d* = .15). There was no significant interaction of SESSION x DIRECTION, *χ^2^*(1, *N* = 37) = 0.347, *p* = .55, suggesting that the effect of tACS on IHI was comparable for both directional projections.

**Fig. 4.**
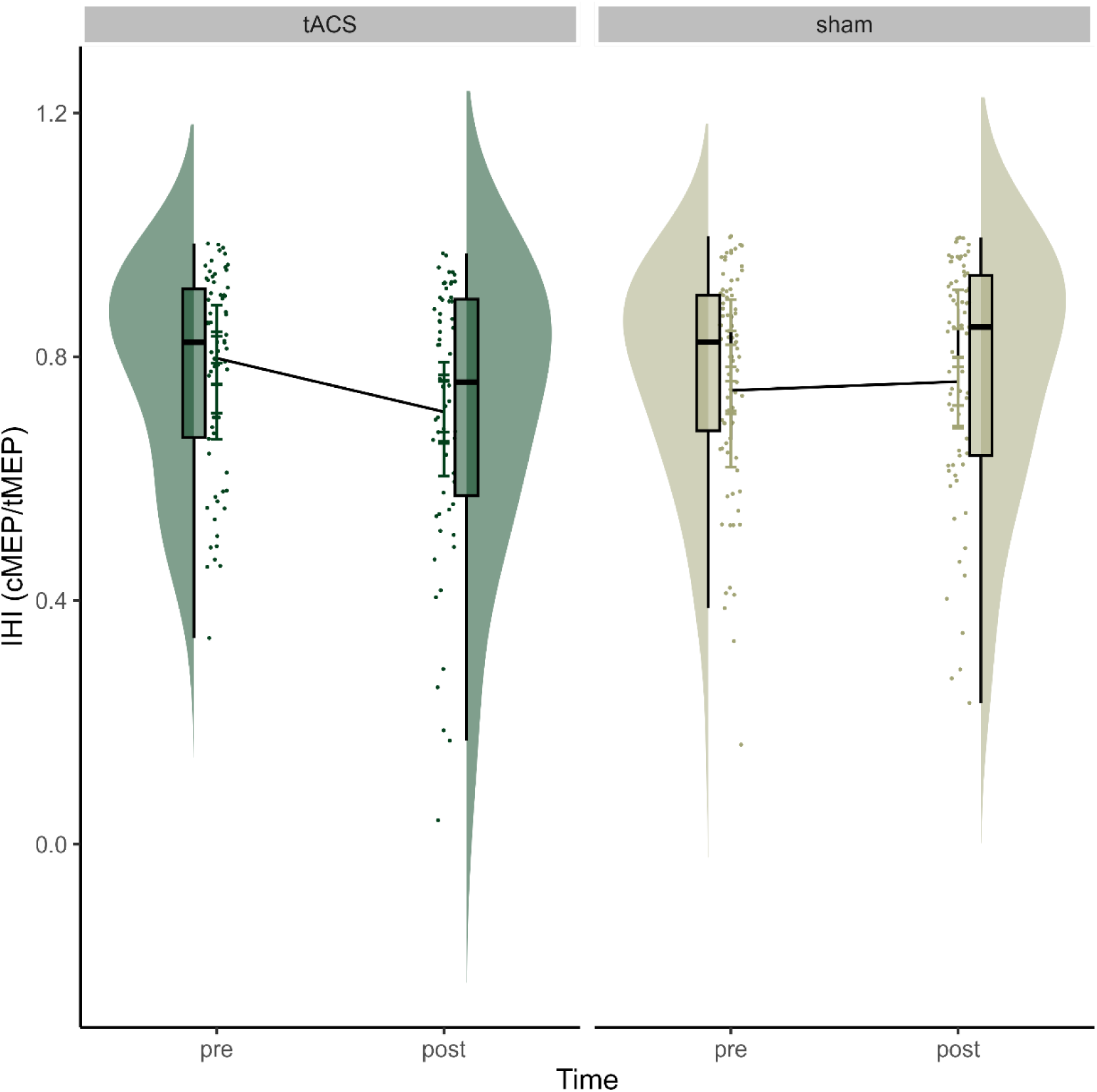
IHI ratio across tACS and sham sessions compared pre-to post-tACS. Half-violins represent the data distribution of IHI ratio at each assessment time point for each session. The boxplots show the median and interquartile range to visually summarise central tendency and dispersion. The horizontal line within the box plot represents the median. The top and the bottom lines represent the upper and lower quartiles, respectively. The data points outside of the whiskers are >1.5 quartiles. The line connecting the paired violins visualizes the direction of IHI changes between pre- and post-stimulation.

### EEG reveals beta power decreases following ds-tACS

For the spectral power analysis of beta-power (20 Hz) changes at electrodes C3 (left M1) and C4 (right M1) following ds-tACS, a GLMM revealed a significant interaction of SESSION x TIME, x HEMISPHERE *χ^2^*(1, *N* = 37) = 4.41, *p =* .04. Follow up contrasts revealed a decrease in beta power at post-stimulation irrespective of the sessions in the left, *z* = −2.71, *p <* .01, *d* = −.39, and right M1, *z* = −5.64, *p <* .01, *d* = −.77, except for the right M1 after sham which showed an increase from pre- to post-stimulation, *z* =.51, *p <*.01, *d* =.21. During both tACS and sham sessions, the left M1 showed a decrease in beta power. While both tACS and sham sessions showed significant beta power decrease in the left M1, follow-up contrasts revealed that the magnitudes of decrease in beta power were significantly greater in the tACS session relative to the sham session irrespective of the hemispheres, *z* = −2.71, *p <* .01, *d* = −.37.

### Functional connectivity indexed by Imcoh did not change following ds-tACS

We investigated whether functional connectivity changed by assessing endogenous oscillatory phase synchronisation at 20 Hz following ds-tACS to the bilateral M1 regions during resting-state EEG. Figure 5 illustrates the ImCoh values at pre- and post-stimulation for tACS and sham sessions. For resting-state ImCoh at 20Hz, there was a significant main effect of TIME *χ^2^*(1, *N* = 37) = 5.54, *p*=.02. Follow-up contrasts revealed that across sessions, ImCoh value decreased from pre- to post-stimulation, *z* = −2.66, *p*<.001, *d*=-.26. The main effect of SESSION, *χ^2^*(1, *N* = 37) = 1.48, *p* = .22 and the interaction SESSION x TIME were not significant, *χ^2^*(1, *N* = 37) = 2.86, *p* = .09.

**Fig. 5.**
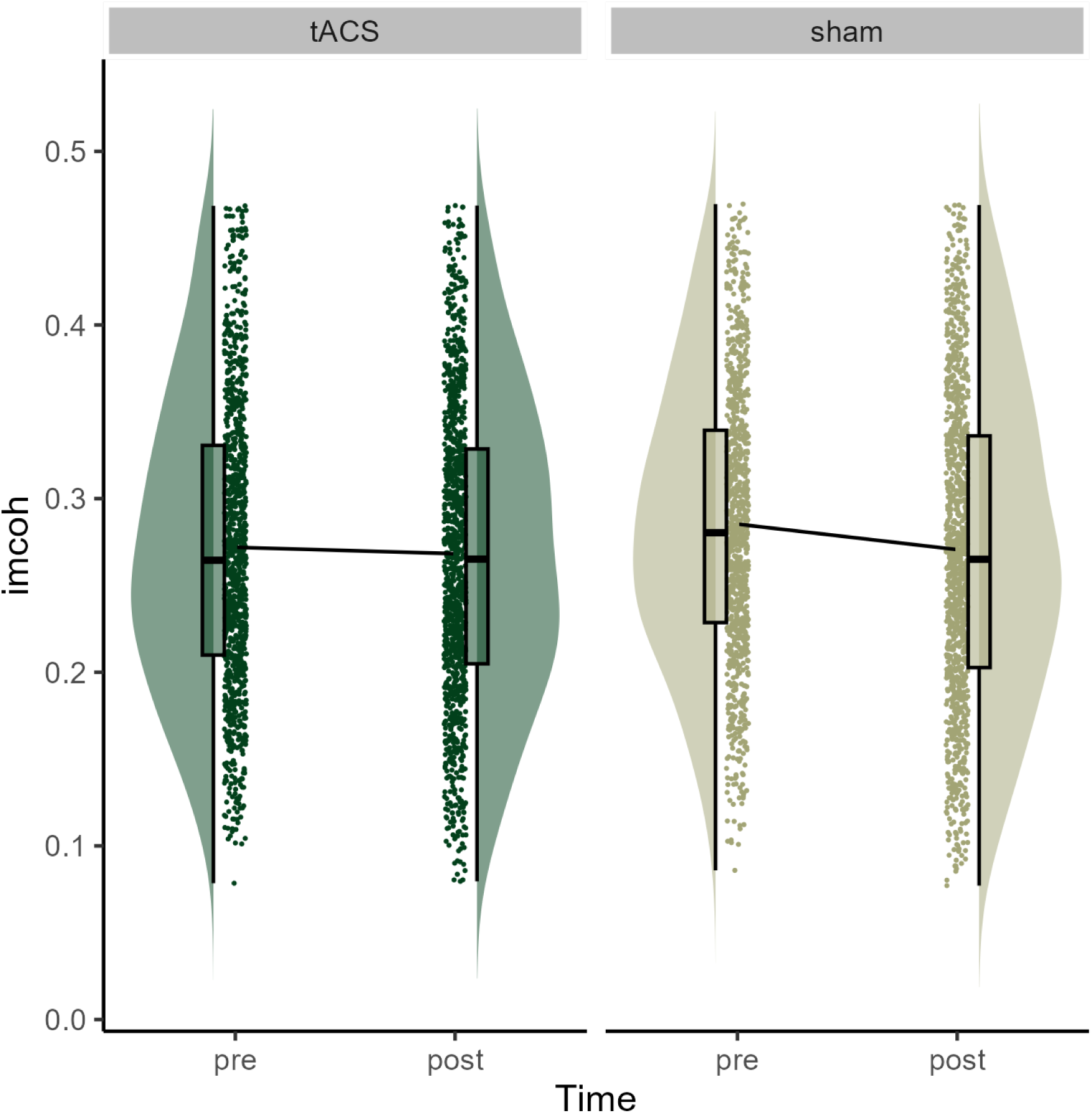
Resting-state imaginary component of coherency (ImCoh) at beta frequency bands across tACS and sham sessions compared pre-to post-tACS. The half-violins represent the data distribution of imCoh at each assessment time point for each session. The width of each violin indicates the density of ImCoh values. The boxplots show the median and interquartile range to visually summarise central tendency and dispersion. The horizontal line within the box plot represents the median. The top and the bottom lines represent the upper and lower quartiles, respectively. The data points outside of the whiskers are >1.5 quartiles. The line connecting the paired violins visualizes the direction of ImCoh changes between pre- and post-stimulation.

## Discussion

The present study sought to elucidate the neural mechanisms that underlie functional connectivity modulation induced by ds-tACS to the bilateral M1s. We found that ds-tACS delivered at 20Hz for 20 minutes to the bilateral M1s induced changes in functional connectivity and behaviour, observed through increased IHI and improved bimanual dexterity. This effect contrasts with the lack of modulation observed in oscillatory-based connectivity measures, highlighting the distinct mechanisms underlying the post-stimulation effects of ds-tACS. The lack of functional connectivity changes in oscillatory measure, i.e., ImCoh, suggests that ds-tACS applied to the bilateral M1s has no impact on post-stimulation measures of oscillatory-based functional connectivity. The observed manual dexterity improvements also indicate that applying ds-tACS over the bilateral M1s might have enhanced the functional connectivity of the circuits underlying bimanual coordination rather than enhancing global motor function, as we did not observe significant changes in the unimanual tasks. The observed increase in IHI indicated that ds-tACS applied over the bilateral M1s specifically enhanced the interhemispheric pathways connecting these distant cortical regions, potentially facilitating the improvement of bimanual coordination.

### Dual-site tACS improved interhemispheric inhibition and corticospinal excitability declined

Our findings showed that IHI increased following beta-tACS to the bilateral M1s, without a corresponding increase in corticospinal excitability but rather a decrease. Previous studies investigating IHI during or following beta frequency tACS application to single-sites in sensorimotor areas found no change in inhibition (Rjosk et al. 2016; Lafleur et al. 2020). Our findings align with previous research, which found that ds-tACS induced changes and enhanced functional connectivity (Violante et al. 2017; Reinhart & Nguyen 2019; Grover et al. 2022; Meng et al. 2023). Beta ds-tACS has also been linked to improvements in response inhibition tasks. Fujiyama and colleagues (2023) found that ds-tACS over the right inferior frontal gyrus and pre-supplementary motor area enhanced beta-band functional connectivity. Thus, in addition to the previous studies finding modulations of EEG-based functional connectivity, our study demonstrated that ds-tACS applied over the bilateral M1 can improve interhemispheric communication, demonstrating connectivity was enhanced between the M1s.

It is important to note that, in the current study, no increases in corticospinal excitability were observed, indicating that our observation of IHI modulation was driven by the specific effect of ds-tACS on interhemispheric pathways rather than an increase in M1 excitability. A meta-analysis by Wischnewski and colleagues (2019) found that single-site beta-frequency tACS consistently increased corticospinal excitability when applied at voltages above 1.0mA. In the present study, dual-site application diverges from this consensus. IHI value can decrease independently of changes in interhemispheric interactions between M1s if corticospinal excitability increases, as the IHI ratio is calculated as MEP_CS-TS_/MEP_TS_. However, we observed a decrease in the IHI ratio despite the overall decline in corticospinal excitability following both tACS and sham sessions. Therefore, the observed decrease in the IHI ratio reflects an increase in interhemispheric inhibition and suggests genuine augmentation in IHI between the M1s. These findings are comparable to Nowak and colleagues (2017) observations that single-site tACS applied over M1 modulated intracortical inhibition with no tACS-related changes in corticospinal excitability. Therefore, our findings suggest that the increase in IHI is not dependent on cortical excitability but instead reflects a targeted effect on interhemispheric pathways, as predicted.

In our investigation of how ds-tACS may modulate different interhemispheric circuits through stimulation, we found no significant differences in modulations between SIHI and LIHI. Interhemispheric inhibitory interactions are mediated through transcallosal pathways and local inhibitory neurons (Daskalakis et al. 2002). It has been suggested that SIHI and LIHI are modulated by GABA_A_ and GABA_B_-ergic circuits, respectively (Irlbacher et al. 2007). Therefore, this lack of significant difference in modulations between SIHI and LIHI implies that while ds-tACs over M1s engages interhemispheric pathways, it does not appear to selectively influence either GABA_A_- and GABA_B_-ergic circuits. Previous studies examining the effect of single-site tACS over M1 on intracortical inhibition found gamma frequency tACS induced greater significant changes in GABA_A_ activity than beta-tACS both during stimulation (Nowak et al. 2017) and following stimulation (Guerra et al. 2018).

Nowak and colleagues (2017) proposed that although beta-frequencies are the predominant oscillations in the sensorimotor cortex, inhibitory neurons may differ and are intrinsically responsive to gamma-band synchrony. Thus, beta-frequency ds-tACS might not be selective in modulating inhibitory pathways mediated by GABA_A_ and GABA_B_-ergic, instead producing a broader or generalised effect across the sensorimotor circuits.

### Dual-site tACS had no impact on oscillatory synchrony

Previous research suggests that ds-tACS is capable of synchronising oscillatory activity across distant cortical regions, as the oscillatory synchronisation of the neural populations in distinct cortical regions is considered a key mechanism for the modulation of functional connectivity (Polanía et al. 2012; Thut et al. 2012). Previous studies found ds-tACS coupled oscillatory activity at two cortical sites post-stimulation at theta frequencies (Polanía et al. 2012; Violante et al. 2017; Reinhart & Nguyen 2019), alpha frequencies (Zaehle et al. 2010; Helfrich et al. 2014) and beta frequencies (Fujiyama et al. 2023; Meng et al. 2023). However, the present results diverged from these observations, as no post-stimulation increases were found in the oscillatory functional connectivity measure (ImCoh). Interestingly, when tACS was applied during task performance (online stimulation), oscillatory increases using the same EEG measure, i.e., ImCoh, were observed (Helfrich et al. 2014; Fujiyama et al. 2023), suggesting that entrainment may primarily underlie online stimulation effects. These findings collectively indicate that beta-frequency ds-tACS applied at rest may induce neuroplastic changes in motor cortical areas through mechanisms distinct from direct oscillatory synchronisation. Another perspective is that the effects of ds-tACS are region-specific and that the heterogeneity of the underlying circuits within sensorimotor areas needs to be considered. Previous evidence indicated phase synchrony was induced by beta-tACs between the right inferior frontal gyrus (rIFG)-pre-supplementary motor area (Meng et al. 2023) and rIFG-M1 (Fujiyama et al. 2023). The rIFG is implicated in response inhibition studies as it is a key component of the inhibitory control network (Aron et al. 2014). Considering beta oscillations are critical for response inhibition (Swann et al. 2009; Wagner et al. 2015), the rIFG may be more susceptible to beta synchronisation than other cortical areas due to its role in response inhibition.

Whilst we found no modulation in ImCoh following ds-tACS, we observed a significant decrease in beta power at post-stimulation, suggesting that ds-tACS interacts with ongoing neural activity. A decrease in beta power has been associated with the activation of the sensorimotor network, particularly in preparation for motor movement (e.g., Kilavik et al. 2013). By integrating these findings with the observed improvement in bimanual dexterity, we may infer that beta frequency tACS may prime cortical areas in a preparatory state for movement as suboptimal preparation likely causes inappropriate execution (Hallett 2000).

Possible explanations as to why beta power showed a decrease following beta-tACS are that the stimulation may have disrupted existing beta oscillations through phase-coupling effects, leading to a reduction in beta power. Additionally, beta-tACS could have modulated excitability in the targeted brain regions, altering the balance between excitatory and inhibitory activity, possibly contributing to the decrease in beta power. The level of beta power can also depend on the difficulty of the task at hand. Bootsma and colleagues (2021) discovered that beta power decreased during challenging motor tasks compared to medium and low-difficulty tasks, indicating a compensatory mechanism to meet increased motor demands. Similarly, Engel and Fries (2010) proposed that in situations where external, bottom-up factors drive behavioural responses, beta power tends to decrease. These findings emphasise the intricacy of relationships between oscillatory activity and task demands. The decrease in beta power following ds-tACS may indicate a shift towards a more adaptable cortical state that supports movement preparation.

While the present results are inconsistent with the expectation that beta power would increase following tACS (Thut et al. 2012), a more nuanced perspective emerges when considering alternative contexts. For example, in people with Parkinson’s disease (PD), reductions in the synchronised activity of beta frequencies during rest were associated with clinical improvements in motor functioning (George et al. 2013). The same study demonstrated that higher beta power in PD patients was associated with more severe motor dysfunction. Similarly, Brittain and colleagues (2014) found that increased beta power in the motor regions of the basal ganglia hindered motor processing in individuals with PD, leading to typical motor impairments such as slowed movements and rigidity. Clinically, reducing this heightened beta activity has been shown to improve symptoms like akinesia and rigidity while lessening tremors (Tinkhauser et al. 2017). However, the present results were derived from a non-clinical population, so we may only speculate on the generalisability of ds-tACS-related power modulations in therapeutic settings.

### Dual-site tACS improved bimanual coordination

We found that bimanual dexterity significantly improved following beta frequency ds-tACS over bilateral M1. The current study is the first to assess manual dexterity in relation to beta ds-tACS. The observed improvement in behaviour following ds-tACS aligns largely with previous studies. For example, Meng and colleagues (2023) found that ds-tACS at beta frequencies targeting the right inferior frontal gyrus and M1 enhanced response inhibition performance. The authors suggested that this improvement in behaviours reflected the neuroplastic reorganisation of brain networks, indicating that, as suggested by the effect of ds-tACS on functional connectivity, it can be broadly transmitted throughout the brain to modulate behaviour.

Improvement in dexterity was only observed in the bimanual assembly task, indicating that improvement was unrelated to the shared properties involved in both unimanual and bimanual assembly, including individual hand movement speed or acuity, but rather a performance improvement in the coordination between both hands. The complexity of bimanual tasks, which necessitate effective communication and coordination between brain regions, may be susceptible to the effect of tACS. These higher-level tasks rely heavily on efficient neural communication and regional coordination (Schoenfeld et al. 2021), suggesting that tACS - which has been demonstrated to enhance connectivity between brain regions - would be more beneficial for such tasks (Schoenfeld et al. 2021). Furthermore, bimanual tasks engage both M1s more intensively than unimanual tasks, potentially fostering greater neural plasticity due to the increased effort required for movement coordination (Feurra et al. 2011). The behavioural task implemented in the current study was not a learning task, as demonstrated by Schoenfeld and colleagues (2021); as such we can only suggest that greater plastic effects may arise from the interhemispheric connectivity changes induced by ds-tACS over bilateral M1s. Consequently, the substantial improvement in bimanual motor performance observed in the current study highlights the potential of M1 beta frequency ds-tACS to enhance bimanual coordination by improving interhemispheric connectivity.

### Limitations and future perspectives

While this study has offered valuable insights into the mechanism underlying ds-tACS, certain limitations must be considered. Individual variability is crucial in discussions of cortical stimulation as it is scientifically unsound to assume that all brains are identical in function and structure (Wansbrough et al. 2024). As such, it may explain the lack of changes in ImCoh following tACS. To minimise individual variability, employing a closed-loop approach is an emerging trend in tES, which involves dynamically adjusting stimulation parameters based on real-time brain activity rather than predetermining parameters (Stecher et al. 2021; Wansbrough et al. 2024). For example, when determining electrode placement or stimulation voltage, structural or functional magnetic resonance imaging (MRI) can be incorporated to locate the optimal electrode placement for the target cortical area (Jog et al. 2021). Individualising tES frequency is another way to tailor brain stimulation. Frequency ranges are well-established as default within specific cortical regions, and when stimulation is adjusted to an individual peak neural oscillatory frequency, neurophysiological changes have been observed (Salamanca-Giron et al. 2021; Aktürk et al. 2022; Kudo et al. 2022; Riddle et al. 2022). Tailoring stimulation creates individualised tES protocols that are proposed as the most appropriate method for ensuring consistency across age, genetics, and existing neural networks and increases the reliability of tES interventions (Wansbrough et al. 2024). In the current study, we employed e-field modelling using a standard head model, which indicated that the protocol used effectively stimulated the M1s with an adequate electrical field (refer to Figure 1A). Future research would benefit from developing individualised protocols that utilise individual participants’ structural MRI scans to fully harness the potential of ds-tACS.

The temporal constraints of our assessment methodology have limited the scope of our findings, specifically the timing and sequencing of assessments before and after stimulation, as shown in Figure 1B. Post-stimulation assessments were conducted between 10-20 minutes of reverse sequencing from the pre-stimulation assessments. Conducting testing this way limited the potential to observe oscillatory changes that may take longer to manifest, but we also lacked insights into the lasting implications of ds-tACS. We included a 7-day washout period to account for lasting effects; however, no follow-up assessments were conducted to determine how long the effects of ds-tACS induced lasted. Research has shown that the effects of tES persist beyond application, with alpha-tACS effects lasting up to 70 minutes (Kasten & Herrmann 2017). Understanding the lasting effects of tACS will inform the longevity of the induced neuroplastic or behavioural changes and provide valuable insight into how tACS is therapeutically beneficial in clinical settings.

Lastly, while the current study was sufficiently powered to detect pre- and post-stimulation changes in neurophysiological measures, future studies are warranted to increase the number of participants. Our exploratory analysis examining the relationship between changes in the IHI ratio and changes in bimanual performance in the PPT revealed a small, non-significant positive correlation (Pearson’s *r* = 0.292, 95% CI [-0.08, 0.59]). Although this correlation was not statistically significant, it does not rule out the possibility of a relationship between modulation in IHI and improvements in bimanual performance in the PPT. Future studies could facilitate correlational analyses to investigate further whether changes in bimanual performance are associated with alterations in interhemispheric inhibition following ds-tACS.

### Conclusion

The current study investigated the neurophysiological mechanisms by which concurrently applied tACS induces changes in functional connectivity between the bilateral M1s. Our findings indicated that tACS enhanced functional connectivity between M1s through IHI, likely through neuroplasticity mechanisms rather than lasting entrainment effects. Further, we observed improvements in bimanual coordination while unimanual dexterity remained unchanged. These findings highlighted tACS as a potential therapeutic tool for neurological conditions marked by dysfunctional cortical connectivity, extending its application beyond the motor system to other cortical regions. Future research may benefit from exploring individualised tACS protocols to fully optimise the effectiveness of ds-tACS. Such effort would facilitate the development of intervention protocols for clinical populations such as PD.

